# Measuring mobility in chromatin by intensity sorted FCS

**DOI:** 10.1101/452540

**Authors:** M. Di Bona, M. A Mancini, D. Mazza, G. Vicidomini, A. Diaspro, L. Lanzanò

## Abstract

The architectural organization of chromatin can play an important role in genome regulation by affecting the mobility of molecules within its surroundings via binding interactions and molecular crowding. The diffusion of molecules at specific locations in the nucleus can be studied by Fluorescence Correlation Spectroscopy (FCS), a well-established technique based on the analysis of fluorescence intensity fluctuations detected in a confocal observation volume. However, detecting subtle variations of mobility between different chromatin regions remains challenging with currently-available FCS methods.

Here we introduce a method that samples multiple positions by slowly scanning the FCS observation volume across the nucleus. Analyzing the data in short time segments, we preserve the high temporal resolution of single-point FCS while probing different nuclear regions in the same cell. Using the intensity level of the probe (or a DNA marker) as a reference, we efficiently sort the FCS segments into different populations and obtain average correlation functions that are associated to different chromatin regions. This sorting and averaging strategy renders the method statistically robust while preserving the observation of intranuclear variations of mobility.

Using this approach, we quantified diffusion of monomeric GFP in high versus low chromatin density regions. We found that GFP mobility was reduced in heterochromatin, especially within perinucleolar heterochromatin. Moreover, we found that modulation of chromatin compaction by ATP depletion, or treatment with solution of different osmolarity, differentially-affected the ratio of diffusion in both regions. Then, we used the approach to probe the mobility of estrogen receptor-α (ER) in the vicinity of an integrated multicopy prolactin gene array. Finally, we discussed the coupling of this method with stimulated emission depletion (STED)-FCS, for performing FCS at sub-diffraction spatial scales.

## Introduction

Chromatin is a macromolecule mainly composed by DNA and histones. Chromatin not only has the function of compacting the DNA in order to make it fit into the nucleus, but also plays an active role in the regulation of all biological processes using DNA as template in eukaryotes, such as transcription, DNA replication and DNA repair. The spatial and temporal organization of chromatin is often deeply perturbed in diseases such as cancer, leading to mis-regulation of these processes and, for example, to aberrant gene expression profiles. From a microscopic point of view, transcription requires the coordination in time and space of multiple macromolecular complexes so that they can timely assemble over an accessible DNA responsive element. Moreover, early experiments clearly established that transcription factors and other nuclear proteins interactions were more dynamic than expected (1, 2). Thus, it remains fundamentally important to determine how proteins move within different regions of the nucleus that are comprised of markedly-heterogeneous chromatin density, and maintain their ability to reach and bind to their target sequences (3).

In this field, a critical role has been played by fluorescence microscopy methods developed to study molecular mobility, including Fluorescence Recovery After Photobleaching (FRAP) (4), Fluorescence Correlation Spectroscopy (FCS) (5) and Single Molecule Tracking (6). These techniques have made it possible to investigate dynamic processes within nuclei of living cells, retrieving information about chromatin dynamics, structure and interactions (7–11). In particular, FCS is based on the analysis of fluorescence intensity fluctuations arising from the passage of fluorescent molecules through a small observation volume (~1 fL). The average amplitude and duration of the fluctuations, extracted from the autocorrelation function (ACF) of the intensity signal, provide information on the concentration and the mobility of the fluorescent particles. FCS is typically implemented in confocal microscopes and possesses a high temporal resolution and the sensitivity of a single molecule method, although without the constraint of having only a few molecules labelled in the field of view; moreover, it induces less photodamage to the cells than perturbation methods (12, 13). For these reasons, FCS has been widely used for measurements in the nucleus, and has been shown to be useful for retrieving information on the nuclear environment in an indirect way (i.e., using an inert probe), for studying the motion of molecules interacting with chromatin and for measuring the mobility of chromatin itself (8, 14–20). In this approach, both the structural and dynamic aspects of the nucleus biology can be investigated, providing a more integrated view of nuclear structure and function.

It is important to determine if variations in the architectural organization of chromatin have a significant impact on the diffusion of surrounding molecules. To answer this question, and to characterize protein mobility in different chromatin regions, several strategies have been proposed that add spatial information to the single-point FCS measurement. Maps of diffusion coefficients have been obtained, for instance, by sequential acquisition of single-point FCS measurements (21, 22), by light-sheet illumination (23, 24) or by parallel acquisition of FCS data at multiple observation volumes (25, 26). Another method that has been used to measure fluctuations at, and between, different points in the nucleus is scanning FCS, which is easily implemented on confocal laser scanning microscopes, but whose temporal resolution is typically limited, by scanning, to the millisecond range (27, 28). Finally, diffusion maps have been recently obtained from the analysis of raster image correlation spectroscopy (RICS) data (29). However, these methods will only provide an accurate description of the mobility properties in different nuclear regions if they are immobile during FCS data acquisition.

A significant advantage can be gained if the different chromatin regions are identified with the help of a reference marker (30). For instance, the intensity of a fluorescent protein can be used to identify specific sub-nuclear regions or, simply, the intensity of a DNA marker (e.g. Hoechst) can be used to identify regions of different chromatin density (31). In this case, the reference intensity can be used as a bona fide marker to assign, during data analysis, each single-point FCS measurement to a specific chromatin region. In this respect, the acquisition of a brief FCS measurement is fundamental to ensure that the probed region does not move significantly during each measurement. On the other hand, the poor statistics resulting from a short FCS acquisition must be compensated by averaging over many FCS measurements assigned to the same chromatin region.

Here, we implemented this idea by performing a slow circular scanning of the excitation beams across the nucleus. The fluctuation analysis is performed by dividing the whole acquisition into a large number of short temporal segments, and considering each segment like an independent FCS measurement, tagged with an intensity value of the reference marker. The ACFs calculated from these short segments are first sorted into two or more populations, corresponding to specific chromatin regions, and then averaged. As a result, this intensity-sorted FCS approach yields, for each measurement, an ACF associated to each chromatin region. In addition, since each measurement is acquired from a single cell, it is possible to measure the dynamic properties of different compartments cell-by-cell, thus avoiding the intracellular mobility differences being distorted due to the intercellular variability.

We validated the technique by measuring differences in the diffusion coefficient of GFP in the nucleolus and in the nucleoplasm of live HeLa cells using, as a reference, the relative intensity variation of GFP in the two compartments. Then, we applied the technique to detect differences of GFP diffusion between regions of hetero- and eu-chromatin, using Hoechst staining of DNA as the reference. We found that mobility of GFP is reduced in the heterochromatin regions, especially in the perinucleolar heterochromatin. This is, to the best of our knowledge, the first time that such a reduction in mobility is observed for a small, inert probe like the monomeric GFP. The ratio between the diffusion coefficient in hetero- versus eu-chromatin was monitored upon treatments affecting chromatin compaction. We found that compaction due to ATP depletion or hyperosmolar treatment affected this ratio in different way. In addition, we measured the mobility of the estrogen receptor-α (ER) on or away from an engineered transcription locus. Finally, we showed that the approach can be combined with stimulated emission depletion (STED)-FCS to obtain sub-diffraction spot-variation FCS data in specific nuclear regions.

## Materials and Methods

### Implementation of intensity sorted FCS

The schematic implementation of the method is depicted in Fig.1a-c. The excitation volume of a confocal microscope is slowly scanned along a circular line across the specimen (Fig.1a), similarly to what is done in orbital scanning (32, 33), while the fluorescence intensity from one or more spectral channels is continuously recorded at high temporal resolution (Fig.1b). The whole intensity trace *I* recorded in one channel is divided into short temporal sequences, or segments, of duration T_seg_ and from each segment a short-sequence (ss) ACF is calculated (Fig.1b). Then the ssACFs are sorted based on the value of intensity *I*_s_ associated to each segment (Fig.1c). The intensity *I*_s_ used for sorting the ACFs can be the intensity recorded in the same or in another channel. For instance, if the orbit is scanned across two distinct regions (Fig.1a), detectable by a difference in the intensity *I*_s_ (Fig.1b), one can obtain an ACF associated to each region by averaging only the ssACFs corresponding to segments whose intensity is below or above a given threshold, respectively (Fig.1c).

**Figure 1.**
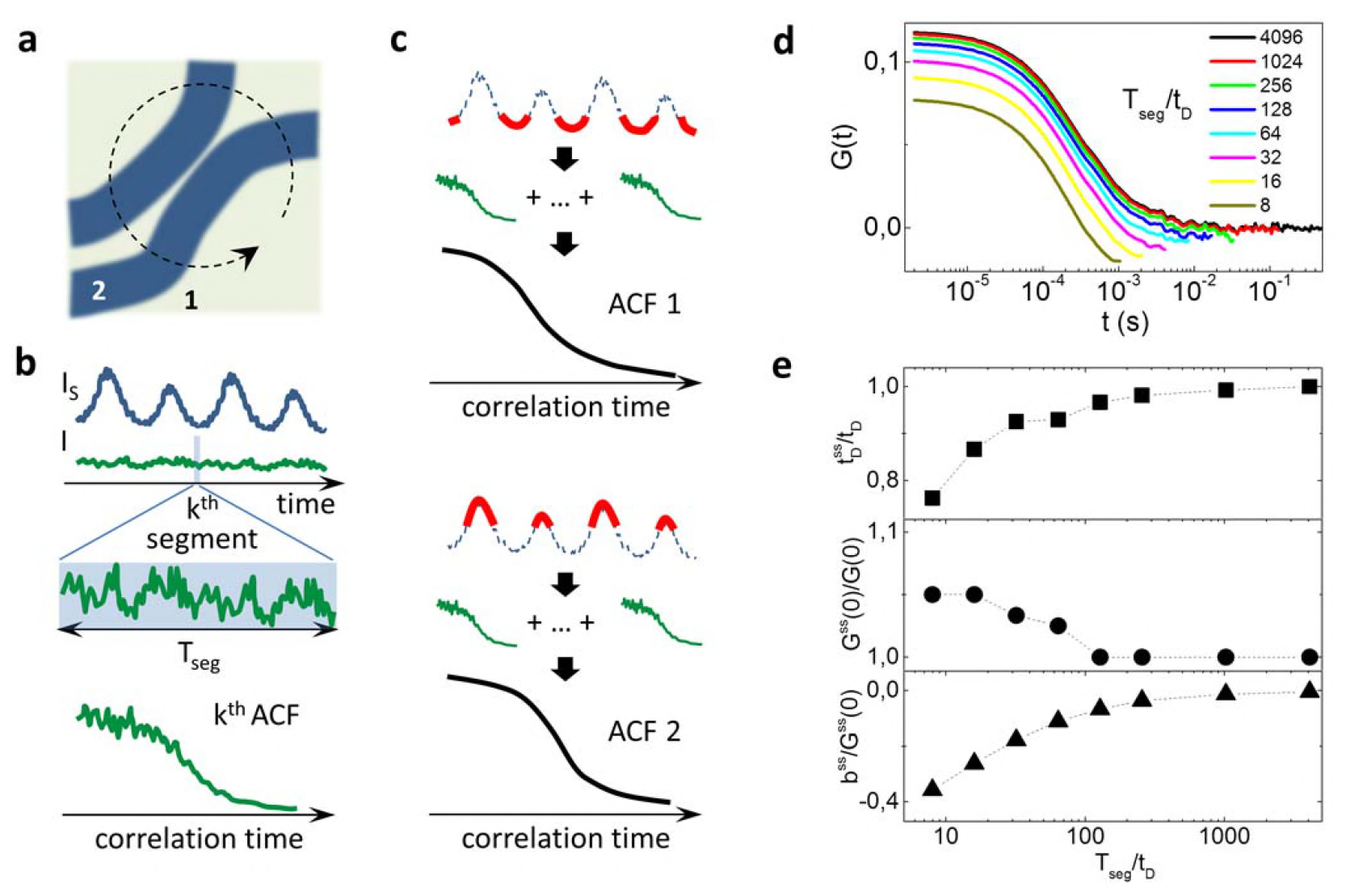
(a-c) Schematic implementation of the method. The excitation spot is scanned in a circular line across the specimen (a) and the fluorescence from one or more channels is continuously recorded (b). The whole measurement is divided into short segments of duration T_seg_ and for each segment the corresponding ACF is calculated (b). The ACFs are then sorted into two or more populations, based upon the value of the reference intensity I_s_, and for each population the average ACF is calculated (c). (d) Short-sequence (ss)ACFs calculated from simulated data for different values of T_seg_/t_D_. (e) Deviation of the fitting parameters (Eq.1) of the ssACF for different values of T_seg_/t_D_.

The duration T_seg_ of the segments must be short enough to resolve intensity variations in the sorting intensity channel *I*_s_ but long enough to properly sample fluorescence intensity fluctuations and prevent deformation of the ACFs. In order to estimate a reasonable lower limit for T_seg_ we simulated molecules undergoing Brownian motion with diffusion coefficient *D* through a Gaussian observation volume of lateral waist *w* and axial waist *w*_z_≫*w*. We divided the resulting intensity trace in segments of duration T_seg_, then the ssACF was calculated and averaged over all segments. The extent of deformation of the ACF depends on the ratio between T_seg_ and the characteristic width of the fluctuations, *t*_D_=*w*^2^/4*D* (Fig.1d). The deformations of the undersampled ACF were quantified by fitting each ACF to the following function:

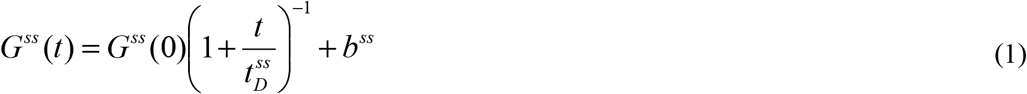

And comparing the fitting parameters with those of the ACF calculated with an infinite sampling time:

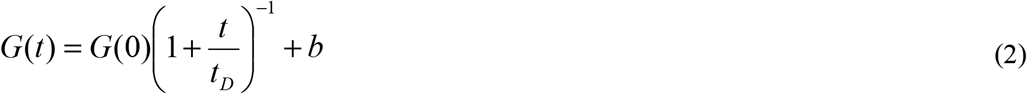

Where *b* is set to 0. For shorter values of T_seg_ the ACF are characterized by shorter values of *t*_D_^s^, negative values of *b*^s^, and slightly larger values of the amplitude *G*^s^(0). We can consider T_seg_∼10^2^*t*_D_ as a lower limit for the duration of the short-sequence. In fact, for T_seg_>10^2^*t*_D_, the more meaningful parameters *t*_D_ and G(0) deviate by less than 5%, in keeping with the rule of thumb that the acquisition time of FCS data has to be at least two orders of magnitude longer than the characteristic correlation time (34).

Another point to take into account is how slow must we scan in order to ignore the correlations due to the motion of the beam. The complete ACF function that describes a model of free diffusion in circular scanning FCS is given by (12):

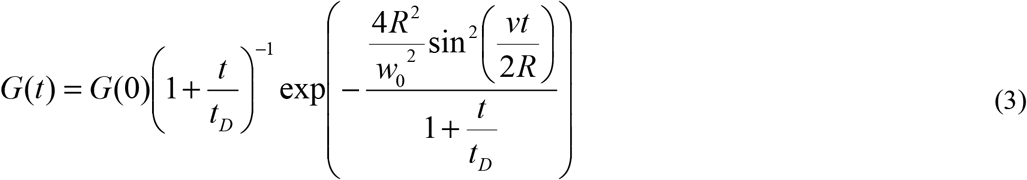

Where *R* is the radius of the orbit and *v* is the scanning speed, given by *v*=2π*R*/*T*, where *T* is the period of the orbit. In our case, since we sample many segments along the orbit, is *t*<*T*_seg_≪*T*, so we can rewrite Eq.(3) as:

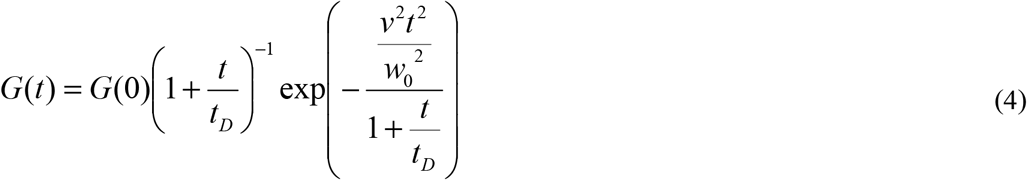

We can ignore the correlations due to the motion of the scanner whenever *v*^2^*t*^2^/*w*_0_^2^≪1+*t*/*t*_D_, namely when *v*^2^≪*w*_0_^2^/*t*^2^+*w*_0_^2^/(*t*_D_*t*). This condition is certainly satisfied if *v*≪*w*_0_/*T*_seg_ and *v*^2^≪*w*_0_^2^/(*t*_D_*T*_seg_). Assuming *T*_seg_=10^2^*t*_D_, the latter condition can also be written as *v*≪0.4D/*w*_0_. For instance, for D≈20μm^2^/s and *w*_0_≈200nm, *v*≪40μm/s. For an orbit diameter of ≈3μm this corresponds to a scanning frequency *f*≪4Hz.

### Simulations

All the simulations were performed using SimFCS (available at http://www.lfd.uci.edu/).

### Optical Setup

The measurements were performed on a custom microscope, obtained from modification of a previous setup (35). The excitation at 485nm was provided by a pulsed laser line (LDH-D-C-485, 80MHz, PicoQuant, Berlin, DE), while the excitation at 407nm was provided by a continuous-wave (CW) diode laser (Cube 1069413/AQ407nm/50mW, Coherent, Santa Clara, CA, US). The STED beam was generated by a CW optical pumped semiconductor laser (OPSL) emitting at 577 nm (Genesis CX STM-2000, Coherent, Santa Clara, CA, US). The laser power was measured at the objective back aperture.

The beams were combined and co-aligned using different laser beam dichroic mirrors, then deflected by two galvanometric scanning mirrors (6215HM40B, CTI, Cambridge Technologies, Bedford, MA, US) and directed toward the 1.40 NA 100x objective (HCX PL APO 100/1.40/0.70 Oil, Leica Microsystems, Wetzlar, DE) by the same set of scan and tube lenses as the ones used in a commercial scanning microscope (Leica TCS SP5, Leica Microsystems, Wetzlar, DE). The fluorescence light was collected by the same objective lens, de-scanned, passed through the laser beam dichroic mirrors, then separated by a fluorescence beam splitter in two channels (detection bands 525/50 nm and 445/45nm) before being focused (focal length 60 mm, AC254-060-AML, Thorlabs, Newton, NJ, US) into fiber pigtailed single-photon avalanche diodes (PDM Series, Micro Photon Devices, Bolzano, Italy). All imaging operations were automated and managed by the software ImSpector (Max Planck Innovation, Munchen, DE) with the exception of circular scanning, managed by the software SimFCS. For FCS measurements, photons were detected by a time-correlated-single-photon-counting (TCSPC) card (SPC-830, Becker & Hickl, Berlin, DE), synchronized with the reference signal provided by the pulsed diode laser.

### Cell Culture

A stable HeLa cell line expressing the protein AcGFP1 was used for all the untagged-GFP experiments (36). The cells were cultured in DMEM supplemented by 10% Fetal Bovine Serum, 2mM glutamine, 100U penicillin and 0,1mg/ml streptomycin (Sigma Aldrich, Saint-Louis, MI, US).

For the estrogen receptor-α (ER) experiments we used a stable HeLa cell line with a hundred-copy integration of the estrogen-responsive unit of the prolactin gene (Sharp et al., 2006), which stably expresses a GFP-tagged version of ER (GFP-ERα:PRL-HeLa cell line) (37). The GFP-ERα:PRL-HeLa cell line was grown in high-glucose DMEM without phenol red supplemented with 5% charcoal dextran stripped Tet(-)FBS, 200ug/ml Hygromycin B and 0.8 ug/ml Blasticidin S (Thermo Fisher Scientific, Waltham, MA, US) and 1 nM Z-4-Hydroxytamoxifen (Sigma Aldrich, Saint-Louis, MI, US).

The day before the experiment, freshly split cells were plated on 8-well chamber (glass bottom, thickness 170±5 µm) (Ibidi, Planegg, DE) and grown overnight.

### Treatments

Nuclear staining was performed incubating the cells for 15min at 37°C with a solution of Hoechst 33342 (Thermo Fisher Scientific, Waltham, MA, US; stock solution 20mM) in PBS, at a final concentration of 4μM. The cells were then washed 4 times with PBS 1X.

Energy depletion was obtained incubating the cells for 30min at 37°C with DMEM supplemented with 50mM 2-deoxyglucose (2-DG, Sigma) and 10mM sodium azide (Sigma Aldrich, Saint-Louis, MI, US) (38). Cells were then imaged directly in ATP-depletion medium.

Treatment with solutions of different osmolarity was performed by addition to the cell of hypo-(190 mOsm) or hyper-osmolar (570 mOsm) solutions for 15min at 37°C (39): the cells were then imaged directly in the incubation solution.

The Sheila cells were treated with 10nM 17-β-estradiol (E2, Sigma Aldrich, Saint-Louis, MI, US) diluted in Live Cell Imaging Solution (Thermo Fisher Scientific, Waltham, MA, US) for one hour to trigger the GFP-ER binding to the array, and then imaged directly in the same incubation solution.

In all the other cases measurements were performed on cells kept in Live Cell Imaging solution.

### Experiments

All the measurements were performed by scanning a circular orbit through the cells nuclei, chosen in such a way to cross the nuclear regions of interest.

For measurements on untagged GFP, the parameters were the following: the 488nm laser power was set to 15μW, while the 405nm laser power at 2,5μW; the orbit diameter was set at 3μm, while the scan period was about 16,7sec. Each measurement was recorded for 132sec. For the ER experiments, the laser powers of the 488nm and 405 nm were set at 5μW and 1μW, respectively; the orbit diameter was 1,5μm, while the scan period was set to about 68 sec, and each measurement lasted 264sec.

For the STED-FCS measurements, the 488nm laser power was set to 15μW, while the STED beam intensity (577nm) was kept at 50mW: the measurements were performed with an orbit diameter of 3μm and a scan period of 16,7sec, while the whole measurement lasted 264sec. For calibration of the effective detection volume, single-point STED-FCS was performed on a solution of purified AcGFP1 (Clontech, Mountain View, CA, US) in PBS as described previously (40). The measurements on solution were performed at an excitation power of 22,5μW for a total acquisition time of 100sec.

### Data Analysis and Fitting

Calculation of the intensity sorted ACFs was performed in Matlab. Each measurement file was first divided into segments, whose duration was calculated basing on the probe mobility: for untagged GFP, the segment duration was set at about 131ms, while for the GFP-ER the segment was 1,05sec long. For each segment, an ACF and an intensity value were calculated. The nanosecond temporal information available in the TCSPC file was used to remove the detector afterpulse in the confocal FCS data, using a custom fluorescence lifetime correlation spectroscopy (FLCS) routine (41). For the STED-FCS data, the nanosecond temporal information available in the TCSPC file was used to generate the multiple ACFs corresponding to sub-diffraction effective volumes, as described in (40). ACFs were only calculated for the green channel, as the Hoechst intensity was used only as a reference channel.

Intensity sorting was performed by averaging all the ACFs of segments whose intensity was below and/or above specific threshold values. Variations of the intensity trace due to photobleaching were removed by a non-linear detrend prior to sorting.

In all the experiments with untagged GFP, the ACFs were fitted using a one-component diffusion model (Eq.2). In the experiments with GFP-ER, the ACFs were either fitted using a two-component diffusion model:

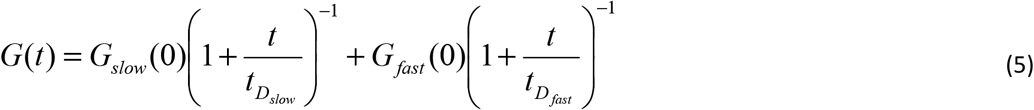

or the diffusion and binding model (Full model) described in (15).

For the two-component diffusion model, a global fit was performed for each experiment, keeping the two values of diffusion coefficients, D_slow_ and D_fast_, shared between the measurements and letting instead vary the amplitudes corresponding to the fast and slow diffusing components.

One- and two-component diffusion fits were performed in Origin. The Full Model fits were performed in Matlab.

## Results and Discussion

### Measurement of the GFP diffusion in the nucleolus vs nucleoplasm

As a validation of the method, we first measured differences in the diffusion coefficient of GFP in the nucleolus and the nucleoplasm of HeLa cells. It has been previously shown that even for a small inert probe like GFP, there is a clear difference in the values of the diffusion coefficient measured in the nucleoplasm in respect to the nucleolus (26, 29, 42). In this case, we used the GFP intensity level as a reference marker to distinguish the two nuclear regions. In fact, the nucleolus of a cell expressing GFP appears dimmer than the nucleoplasm due to a different concentration of GFP in the two compartments (Fig.2 a, b). The intensity trace showed easily-detectable regions of low and high intensity, corresponding to the nucleolus and the nucleoplasm, respectively (Fig. 2 c). By specifically selecting only the short FCS segments corresponding to these low and high intensity regions (Fig. 2 c), we generated the sorted ACFs corresponding to the nucleolus (Fig. 2 e) and the nucleoplasm (Fig. 2 f).

**Figure 2.**
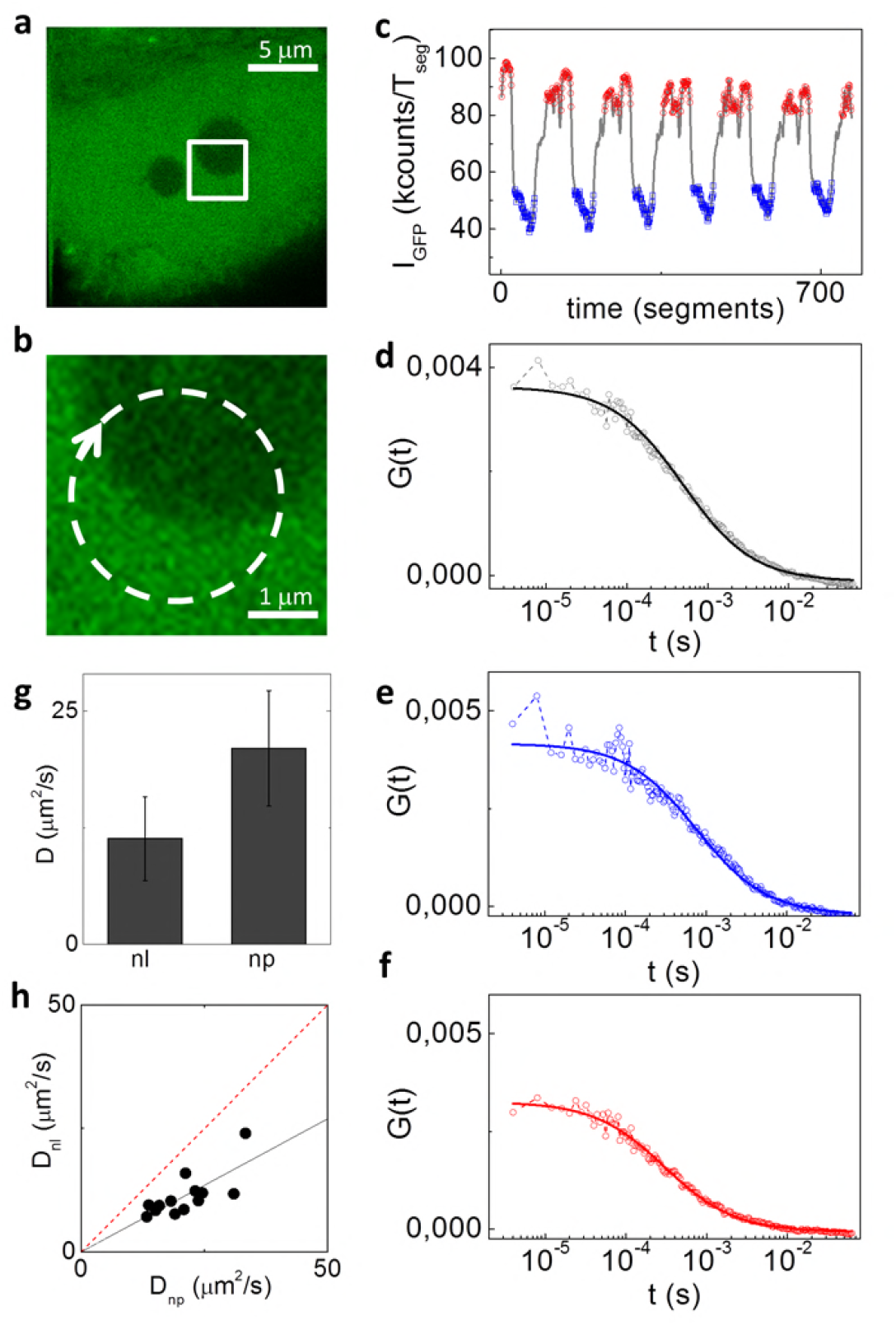
Measurement of GFP diffusion coefficient in the nucleoplasm and the nucleolus. Each measurement is performed on an individual cell (a), scanning the beams across the nucleolus (b). The GFP intensity trace is used for sorting the ACFs between the nucleolus (in blue) and the nucleoplasm (in red) (c). The total ACF without sorting (d) is compared to the mean ACF calculated from the fluctuations measured in the two regions (e, f). The average diffusion coefficient extracted from the fitting of the ACF of the nucleoplasm is significantly higher (24,3 ± 6,2μm^2^/s) than the one shown by GFP in the nucleolus (12,5 ±2,5μm^2^/s) (g). Plot of the ratio between the diffusion coefficients calculated in the two compartments (h), with a linear fit (solid black line, the intercept is fixed to zero) yielding a slope of 0.43. The dashed red line represents the case in which the diffusion coefficients are the same in both the compartments.

By the fit of the ACF we retrieved the averaged (n=13 cells) diffusion coefficient of GFP in the nucleolus (11,3 ±4,5μm^2^/s, mean ± s.d.) and in the nucleoplasm (21 ± 6,2μm^2^/s) (Fig. 2 g). These values are in keeping with the values reported in literature (26, 29, 42), demonstrating that our analysis method works properly. Interestingly, there is a high intercellular variability in the measured absolute values of diffusion coefficient (Fig. 2 g, h). However, the ratio between the two values of diffusion coefficient in each cell is quite conserved, as shown by a D_nl_ vs D_np_ scatter plot (Fig. 2 h). We have evaluated this ratio by performing a linear fit of the data for each independent experiment (Fig. 2 h, Supplementary Fig S1): we obtained D_nl_/D_np_ =0,45 ± 0,02 (mean ± s.d. of 5 independent experiments, Supplementary Fig S1).

These results show, as expected, that the diffusion of GFP is reduced in the nucleolus with respect to the nucleoplasm, due to higher molecular crowding. It is worth noting that we have not used the FCS segments corresponding to the interface between the two regions (Fig.2c), as they are expected to show a mixed behavior. However, the capability of measuring mobility of proteins at the boundary of nuclear domains could be of interest for models of chromatin organization based on phase separation (43).

### Measurement of GFP diffusion in euchromatin vs heterochromatin

Next, we checked if the technique was able to detect differences of GFP diffusion between regions of high and low chromatin density (hereafter referred to as hetero- and eu-chromatin) using as a reference the intensity of Hoechst-stained DNA (Fig. 3). First, we performed the measurements in the nucleoplasm of HeLa cells, in regions far from the nucleolus (Fig. 3 a, b). In this way, we could use the Hoechst intensity as a quantitative reference for nuclear DNA concentration, defining regions of eu-chromatin (low Hoechst signal) and hetero-chromatin (high Hoechst signal) (Fig. 3 c) and to generate the corresponding ACFs (Fig. 3 d, e). We found that there is a slight difference in the values of the diffusion coefficient of GFP (n=23 cells) between euchromatin (21,8 ± 5,6 μm^2^/s) and heterochromatin (19,6 ± 5,4 μm^2^/s), although this difference is not significant when considering the average of measurements performed on multiple cells. Conversely, when we scatter plotted the two values of diffusion coefficients measured in each cell (Fig. 3 f), we found that the ratio between the diffusion coefficient in hetero- and eu-chromatin was less than 1 (D_hc_/D_eu_= 0,87 ± 0,05, mean ± s.d. of 5 independent experiments, Supplementary Fig. S2), despite the high variability between measurements performed on different cells. In fact, the single cell sensibility of our method facilitates stressing the differences in protein mobility within different chromatin regions, without being affected by the intercellular variability.

**Figure 3.**
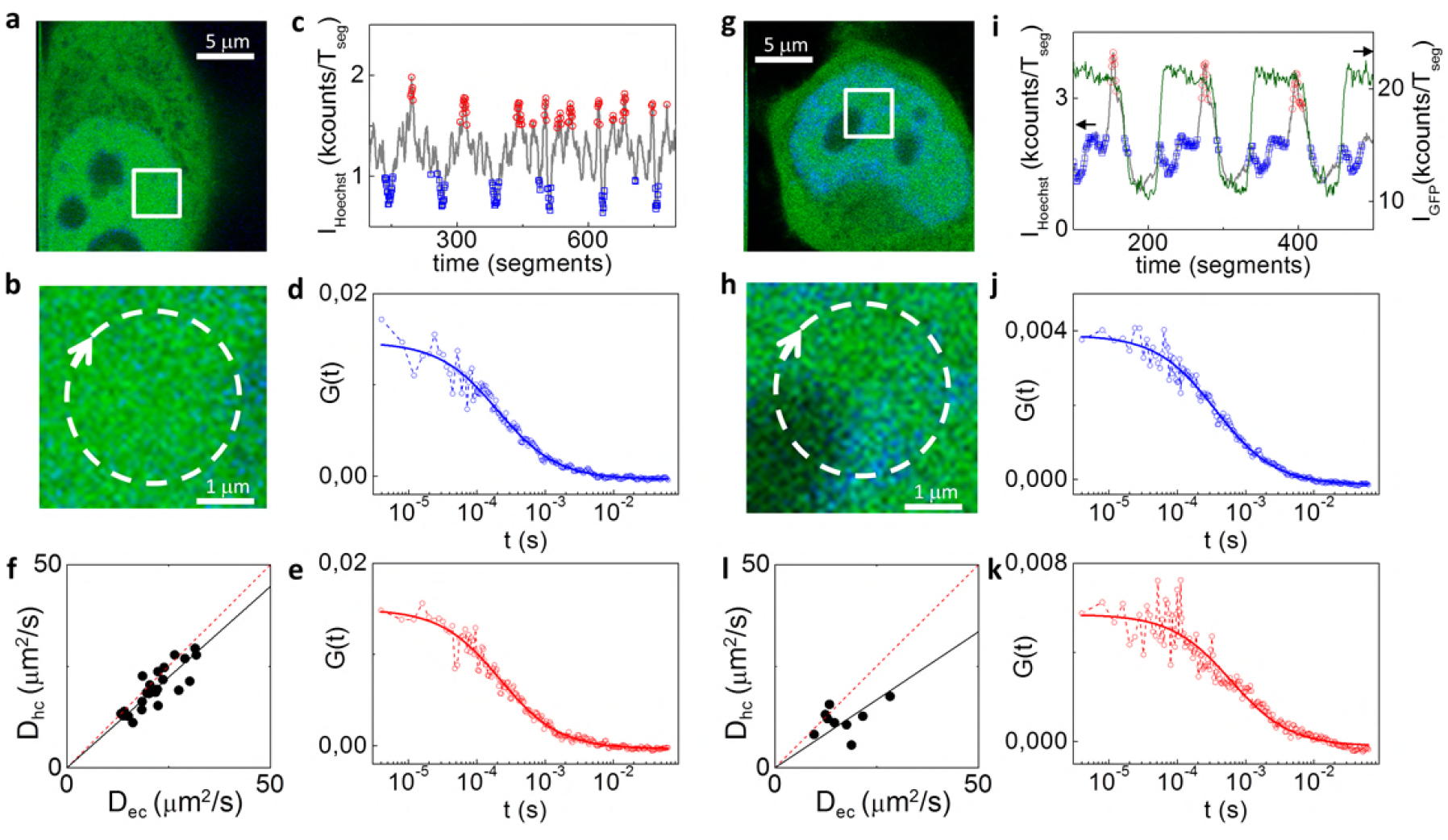
Measurement of the diffusion coefficient of untagged GFP in eu- vs hetero-chromatin (af) and in euchromatin vs perinucleolar heterochromatin (g-k). (a-e) Each measurement is collected from a single nucleus stained with Hoechst (in blue in a, b), far from the nucleolus (b). Basing on the Hoechst intensity trace (c), the FCS segments are sorted into two populations: the heterochromatin, in red, and the euchromatin, in blue. The averaged ACFs are then calculated and a single component fit was performed, giving a value of D_ec_=31,5μm^2^/s for the euchromatin (d) and D_hc_=29,4μm^2^/s for the heterochromatin (e). (f) Scatter plot of the two values of diffusion coefficients D_hc_ and D_eu_ measured on different cells, showing a slightly higher diffusion coefficient in the euchromatin (D_hc_/D_eu_= 0,89). (g-k) When selecting the perinucleolar heterochromatin, the beams are scanned through the periphery of a nucleolus (g,h) and both the GFP (green line) and Hoechst (gray line) intensities are used as references (i). The two average ACFs corresponding to the euchromatin (j) and perinucleolar heterochromatin (k) are calculated and fitted, providing a value of 17,6μm^2^/s and 10,6μm^2^/s, respectively. When results from the analysis of multiple cells are displayed in a scatter plot(l), a more marked difference between the two values of diffusion coefficient is observed (D_hc_/D_eu_ =0,7).

Interestingly, we found a greater difference in the case of perinucleolar heterochromatin (Fig. 3 g-k). To focus on this region, we scanned the beams across the perinucleolar heterochromatin (Fig.3g-h) and used both the GFP and Hoechst intensities as references to discard the FCS segments belonging to the nucleolus (low GFP signal), and generate the ACFs corresponding to the perinucleolar heterochromatin (high GFP signal, high Hoechst signal) and the euchromatin (high GFP signal, low Hoechst signal) (Fig. 3 i-k). For the perinucleolar heterochromatin we obtained a ratio of D_hc_/D_eu_= 0,7 ± 0,07 (mean ± s.d. of 4 independent experiments, Fig. 3 l, Supplementary Fig S3).

Previous FCS studies have been incapable of identifying differences in the mobility of monomeric GFP in nuclear compartments with different chromatin density (21). Our results show instead that even the motion of a small inert probe (monomeric GFP) is affected by the higher degree of compaction of the heterochromatin regions, especially in the perinucleolar heterochromatin. In this respect, we believe that a higher accuracy in our data may result, at least in part, from the following characteristics of our method. First of all, the high (microsecond) temporal resolution of the ACF ensures a proper sampling of the ACF, especially if compared to scanning FCS (millisecond temporal resolution). Second, the efficient sorting of the short FCS measurements ensures that fluctuations are averaged only between regions with the same intensity-based fingerprint (for instance, in the case of heterochromatin, these are only the regions identified by the Hoechst peaks), even if these regions are not completely immobile during the whole acquisition. This is conceptually similar to performing FCS on a tracked subcellular region (44–46), although this tracking is performed *a posterior*i on the recorded intensity profile. A similar idea has been exploited in the context of RICS to perform fluctuation analysis on specific organelles (30). Finally, another advantage of our approach is the possibility of obtaining cell-by-cell estimates of the diffusion coefficients for each of the probed nuclear regions.

### Monitoring the diffusion coefficient of GFP during chromatin compaction changes

In order to test to the extent that chromatin compaction affects GFP diffusion in different chromatin regions, we treated the cells with solutions known to induce changes in the compaction of chromatin.

Solutions of different osmolarities induced visible changes in nuclei morphology (Fig. 4 a, e, i) that are reflected in a large difference in the diffusion coefficients of GFP measured in different compartments. Indeed, if the cells were treated with a hypo-osmolar solution, the diffusion coefficient of GFP was higher than in controls; when the cells were subjected to hyper-osmolar treatment, the diffusion coefficients calculated in both eu- and hetero-chromatin were significantly lower (D_eu_=7,2 ± 2,3 and D_hc_=6,7 ± 2,3 μm^2^/s, mean ± s.d., n = 10 cells) compared to the controls (Table 1).

**Figure 4.**
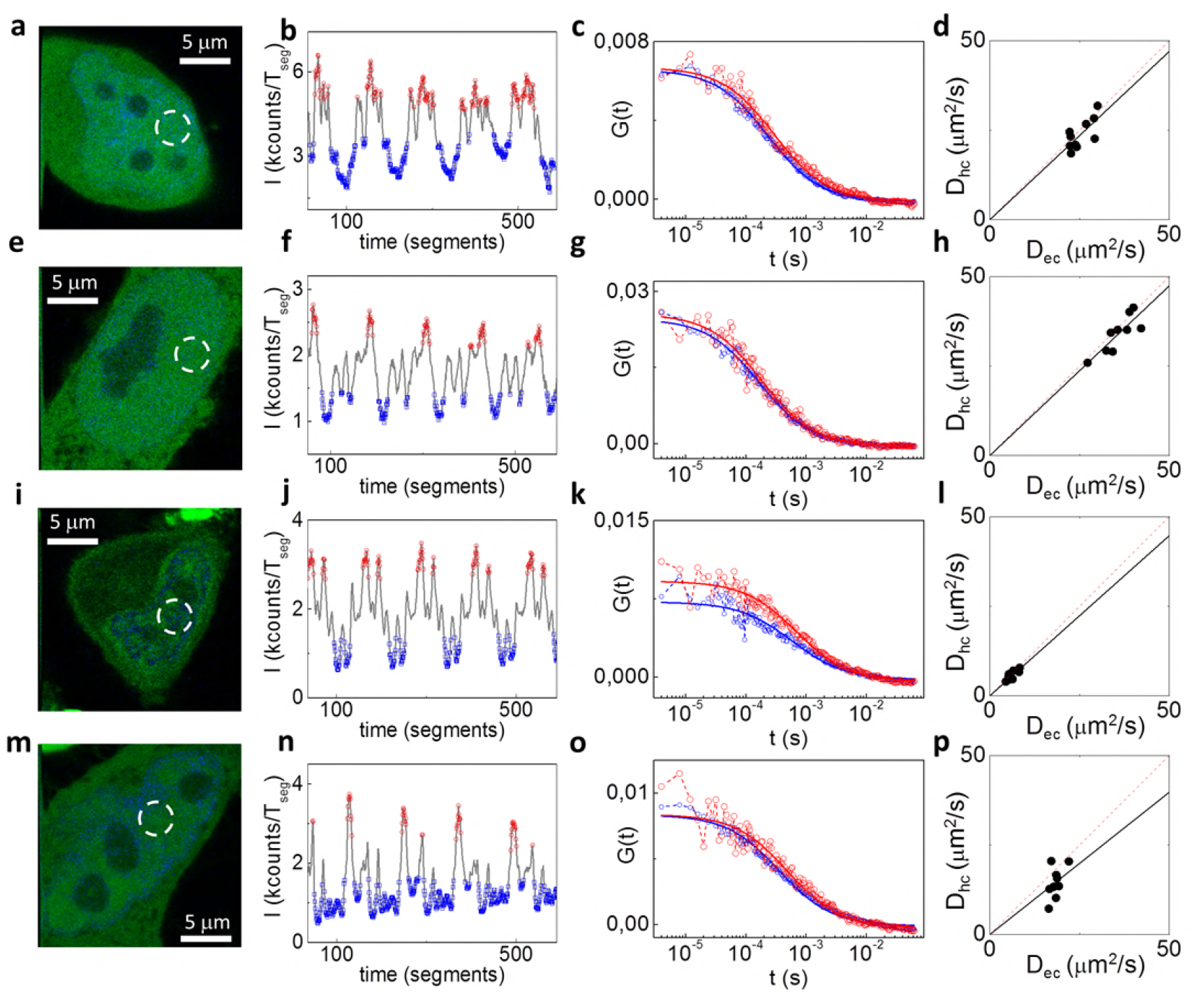
Measurement of the untagged GFP diffusion coefficient after treatments affecting chromatin compaction state. All the measurements were performed comparing eu-and heterochromatin in the nucleoplasm of HeLa cells. (a-d) control cells; (e-h) cells treated with a hypo-osmolar solution; (i-l) cells treated with a hyperosmolar solution; (m-p) cells treated with an ATP-depletion solution. The FCS segments are sorted based on the Hoechst intensity (b, f, j, n), to obtain average ACFs for eu-(blue) and hetero- (red) chromatin (c, g, k, o). With respect to the control (d, D_hc_/D_eu_=0,94), the osmotic treatment greatly affects the absolute values of the diffusion coefficients, but not the ratio between them (h, D_hc_/D_eu_=0,95; l, D_hc_/D_eu_=0,89). The ATP-depleted cells show a less marked difference in the absolute values, but a higher difference between the diffusion coefficients calculated in the two compartments (p, D_hc_/D_eu_=0,79).

**Table 1.** Values of the average diffusion coefficient of untagged GFP in eu- (D_eu_) and hetero-chromatin (D_hc_) and average value of the ratio D_hc_/D_eu_.

| | $D_{eu} (\mu m^2/s)$ | $D_{hc} (\mu m^2/s)$ | $D_{hc}/D_{eu}$ |
| --- | --- | --- | --- |
| Control | $23,5 \pm 8,1$ | $19,4 \pm 6,1$ | $0,87 \pm 0,05$ |
| ATP depletion | $18,1 \pm 4,7$ | $15 \pm 4,9$ | $0,79 \pm 0,08$ |
| Hyperosmotic | $7,2 \pm 2,3$ | $6,7 \pm 2,3$ | $0,92 \pm 0,03$ |
| Hyposmotic | $30,1 \pm 7,8$ | $29,6 \pm 6,8$ | $0,98 \pm 0,03$ |

Interestingly, both hypo- and hyper-osmolar treatments affected only the absolute values of the diffusion coefficients, but not their average ratio D_hc_/D_ec_ (Table 1, Fig. 4 h, l), meaning that the treatment has a similar impact on both eu- and hetero-chromatin compartments. Incubation with an ATP depletion solution induced a visible compaction of chromatin with respect to the control (Fig. 4 a, m), which led to a reduction of GFP diffusion coefficients in both eu- and hetero-chromatin (Table 1). In this case, however, the scatter plot of D_hc_ vs D_ec_ (Fig. 4 p) indicates that ATP depletion results in a more prominent slow-down of GFP diffusion in heterochromatin, possibly as a consequence of a larger increase of compaction in hetero- with respect to eu-chromatin.

### Mobility of a transcription factor in different chromatin regions

As a model of a protein interacting with chromatin, we studied the mobility of the estrogen receptor-a (ER), a transcription factor member of the nuclear receptor super-family and involved in regulation of specific genes in response to hormone binding. In particular, we measured differences in the mobility of GFP-ER inside and outside an engineered, readily-visible prolactin reporter gene “array” following stimulation with 10nM 17-β-estradiol for 1 hour (Fig. 5 a, b). Using GFP-ER intensity as a reference, we measured the diffusion inside (high GFP-ER signal) and outside (low GFP-ER signal) the array (Fig. 5c). The intensity-sorted ACFs were fitted using either a two components pure diffusion model or a Full Model (FM) taking into account diffusion and binding (15).

**Figure 5.**
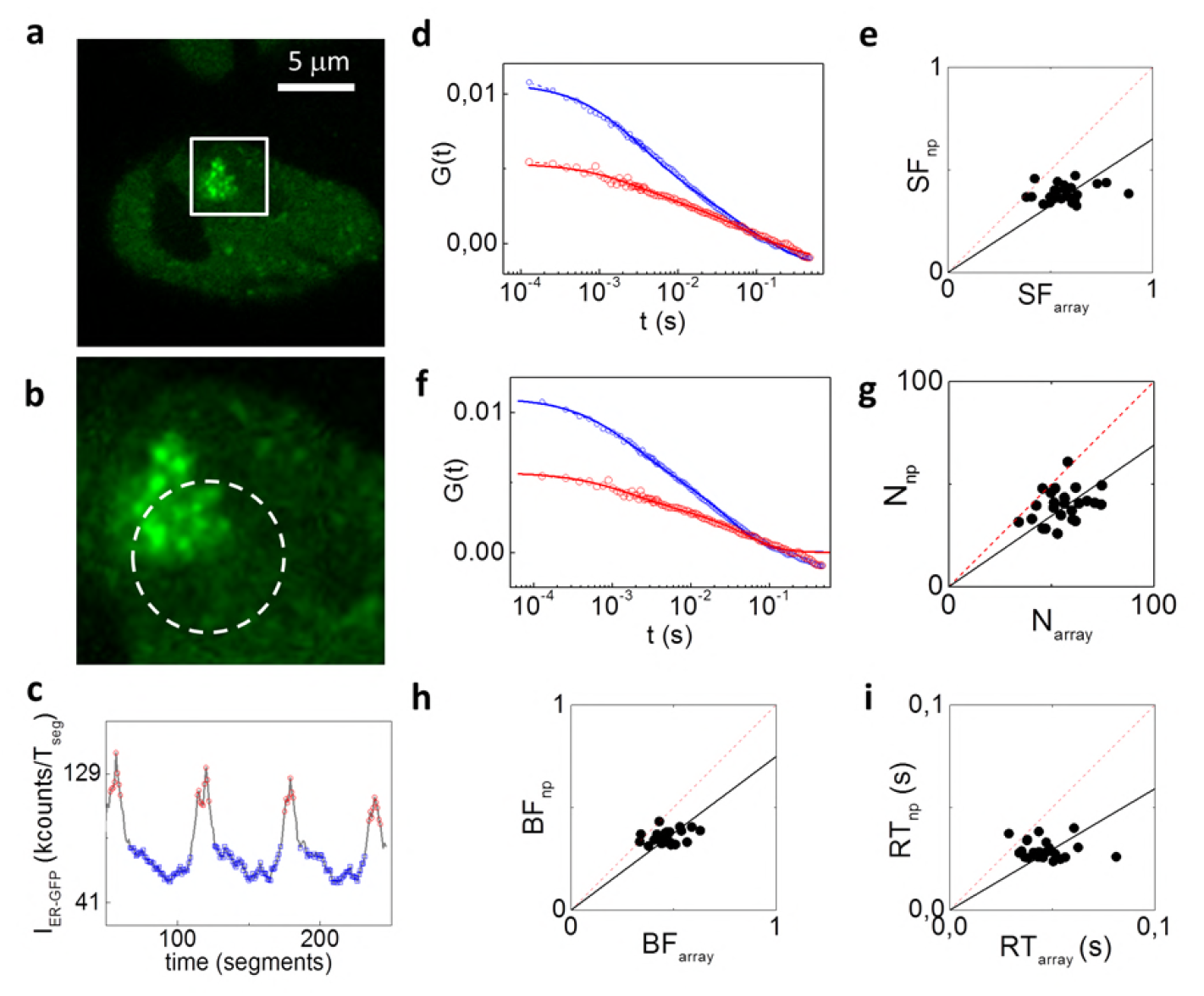
Measurement of the mobility of the estrogen receptor (ER) inside and outside an engineered prolactin gene array. (a, b) The measurements were performed orbiting across the prolactin gene array. (c) The intensity of the GFP-ER signal is used as a reference for sorting the ACFs corresponding to the array (red) and to the nucleoplasm (blue). (d,e) Analysis with a two diffusion component model. (d) A global fit performed on the sorted ACFs (red: sorted ACF of the array, blue: sorted ACF of the nucleoplasm) yields a shared value of diffusion coefficients D_slow_=0,07μm^2^/s for the slow diffusing population and D_fast_=2,1μm^2^/s for the fast diffusing one. (e) Scatter plot of the slow fraction calculated in the two probed nuclear regions (SF_array_/SF_np_= 0,65). (f-i) Analysis with the Full Model, showing the fitted ACFs (f) and the scatter plots of the number of proteins (g, N_np_/N_array_ =0,69), the bound fraction (h, BF_np_/ BF_array_ = 0,75) and the residence time (i, RT_np_/RT_array_= 0,59) for the nucleoplasm and for the array.

In the first case we identified a slow (D_slow_=0,07 μm^2^/s) and a fast diffusing component (D_fast_=2,1 μm^2^/s), in keeping with previous reports (2). We then plotted the slow fraction (SF), calculated as SF=G_0slow_/(G_0slow_+G_0fast_), inside (SF_array_) and outside (SF_np_) the array (Fig. 5 e). As a result of the fit with the two-component diffusion model (Fig. 5 d), we found that the slow diffusing fraction was significantly higher in the array compared to the nucleoplasm, with a ratio SF_array_/SF_np_= 0,73 ± 0,06 (mean ± s.d. of 3 independent experiments, Fig. 5 e and Supplementary Fig S6).

We then performed a fit of the data using the FM model, which is more general and yields several outputs including the number of particles (N), the bound fraction (BF) and the protein residence time (RT) on its binding site (15). The results of the analysis with this model are shown in (Fig. 5 g-i). Reflecting the increased density of estrogen response elements (EREs) at the engineered transcription locus, the number of ER molecules was higher in the array than in the nucleoplasm (N_np_/N_array_=0,67 ± 0,03, mean +/- s.d. of 3 independent experiments, Fig. 5g and Supplementary Fig. S7). Also, the fraction of molecules in a bound state is significantly higher on the array (BF_np_/ BF_array_ =0,83 ± 0,07, mean +/- s.d. of 3 independent experiments, Fig. 5h and Supplementary Fig. S7). Finally, the average time the ER is found in the bound state is longer on the array (RT_np_/RT_array_= 0,65 ± 0,04, mean +/- s.d. of 3 independent experiments, Fig. 5i and Supplementary Fig. S7). These results show that not only is more ER targeted to the gene array (e.g., brighter signal), but also a larger fraction of ER is in a bound state – rather than freely diffusing – at the gene array. Although there are thousands of ER binding sites present throughout the nucleus, ERE density within the prolactin gene array recruits and retains the receptor longer than elsewhere in the nucleoplasm.

### Measurements of GFP diffusion in the nucleolus and the nucleoplasm at sub-diffraction spatial scales

Finally, we tested if the intensity-sorted approach was compatible with super-resolved FCS. Indeed, FCS can be combined with STED microscopy (STED-FCS) to perform fluctuation analysis on sub-diffraction observation volumes (47). In STED, the size of observation volume can be easily tuned by changing the depletion power (48). Thus, an important aspect of STED-FCS is the capability to probe diffusion at different sub-diffraction spatial scales, i.e. to perform a spot-variation FCS analysis, just by changing the depletion power (49). Alternatively, the STED observation volume can be tuned, at a given STED power, by exploiting the fluorescence lifetime variations generated in a CW-STED microscope (50, 51). This strategy has the advantage that a full spot-variation dataset can be obtained in a single measurement, without the need of performing multiple acquisitions at different depletion powers (52). Following this strategy, we recently demonstrated that, by combining the analysis of lifetime variations generated in a CW-STED microscope (51) with FLCS (41), it is possible to perform STED-based spot-variation FCS in single points in the interior of the cell (40).

We coupled this method with intensity sorted-FCS. The measurements were done in HeLa cells, orbiting across the nucleolus and using as a reference the GFP intensity level (Fig. 6 a, b). The GFP lifetime variations induced by the STED laser beam were used to filter the detected photons, identifying in this way three different effective volumes (Supplementary Fig. S8) in the same measurement (40). The sorting of the data generated, for each cell, two sets of ACFs, corresponding to the nucleoplasm (Fig, 6 c) and the nucleolus (Fig, 6 d). This corresponds to a separate spot-variation analysis for each compartment, shown as the average diffusion coefficient versus the square size of the effective observation volume (Fig. 6 e). At smaller spatial scales, we observed a slight increase of the diffusion coefficient, especially in the nucleolus, but also a much larger error bar. This is probably due not only to cell-to-cell variability but also to the fact that reducing the size of the effective detection volume (w_eff_ = 140, 130 and 115 nm, from top to bottom) results in ACFs with a poorer signal-to-noise ratio (Fig, 6 c,d). We tested if the ratio between the diffusion coefficient in the nucleolus versus nucleoplasm was also preserved at different subdiffraction spatial scales, on a cell-by-cell basis (Fig. 6 f). The results are in keeping with those obtained with confocal intensity-sorted FCS (Fig.2), despite a higher scattering of the diffusion coefficients values that are probably ascribed to the SNR reduction of the ACFs at the smallest effective observation volumes.

**Figure 6.**
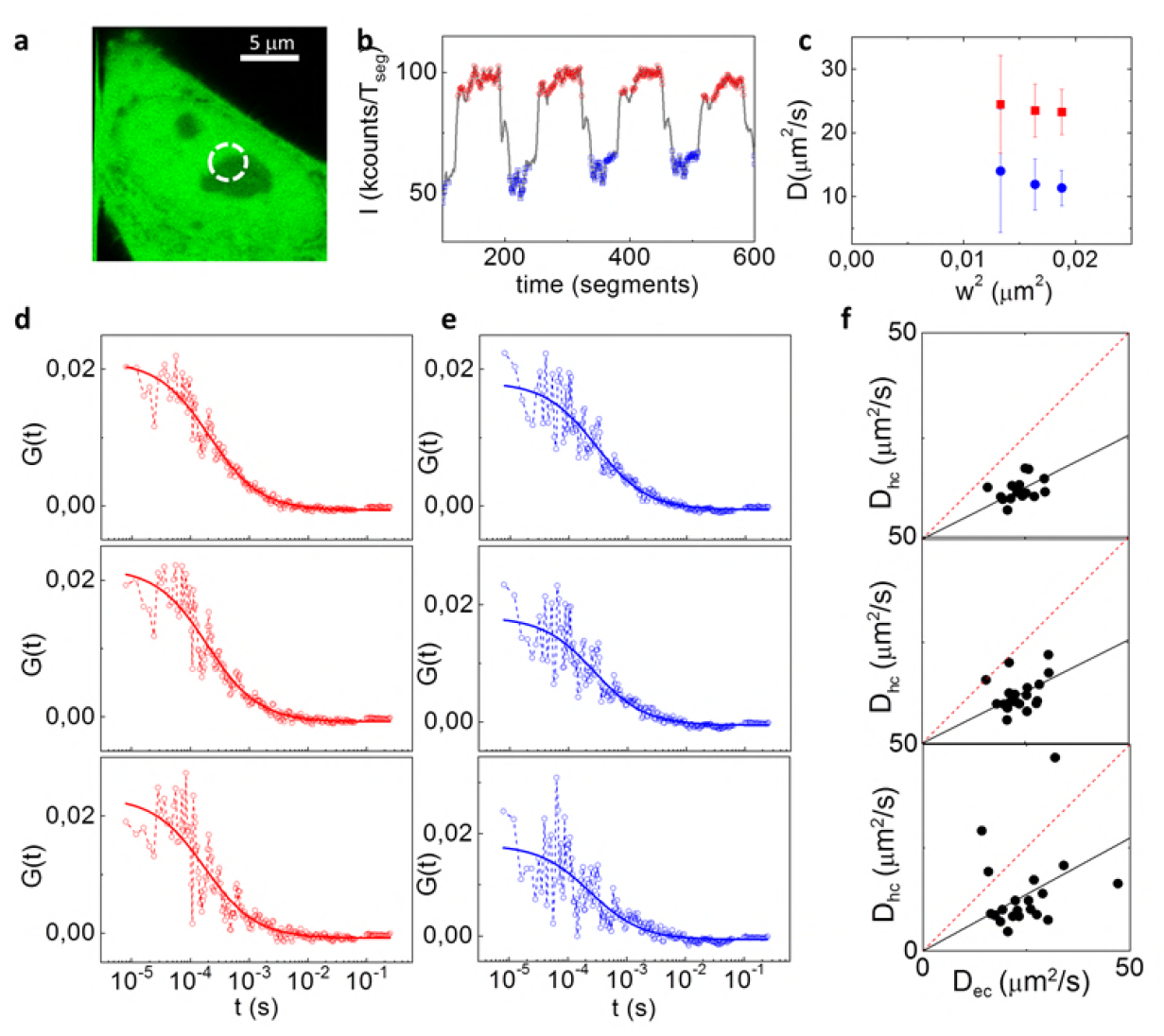
Intensity-sorted STED-FCS in the nucleus. Each measurement is performed on an individual cell, scanning the beams across the nucleoli (a). (b) GFP intensity trace used for sorting the ACFs between the nucleolus (in blue) and the nucleoplasm (red). Three different volumes are selected in the post-processing, with size w=140, 130 and 115 nm respectively. (d,e) ACFs corresponding to detection volumes of decreasing size (from top to bottom) along with the fit, for the nucleoplasm (D_np_=30; 32; 35 μm^2^/s from top to bottom) (c) and the nucleolus (D_nl_=22; 23; 26 μm^2^/s from top to bottom) (d). (e) Average diffusion coefficient versus the square size of effective volume, for GFP in the nucleoplasm (red) and in the nucleolus (blue) (data represent mean ± s.d. of n=20 measurements on different cells). (f) Scatter plot of the diffusion coefficients measured in the two regions for each effective detection volume, together with the corresponding linear fit (D_nl_/D_np_=0.48; 0.5; 0.55 from top to bottom).

Taken together, these results show the compatibility of our approach with STED and that, in principle, intensity-sorted STED-FCS could be a useful tool to measure mobility in different nuclear regions at multiple sub-diffraction spatial scales. Note that the high temporal resolution allows an optimal temporal sampling of the ACF even if the average transit time measured in the nucleoplasm at the smaller effective observation volume is in the order of 100 μs. The major limitation we encountered was related to the poor quality of the ACFs at smaller observation volumes. This aspect could be improved by using probes which are brighter and/or more photostable than GFP, for example using the genetically-encoded Halo-or SNAP- tags, that allow the use of brighter, more stable organic dyes. The increase in the number of photons collected could facilitate pushing the spot-variation analysis towards smaller spatial scales.

Finally, it is worth noting that STED, by providing intensity features that are better resolved spatially, should also improve the sorting of the data. In our test system, i.e. nucleolus vs nucleoplasm, we did not exploit this advantage as the two regions extend over several micrometers and are easily distinguished with normal resolution. Moreover, by averaging all the segments within each compartment, we are prevented from detecting any spatial and temporal heterogeneity within each region. However, this an interesting aspect that might be worth of future investigations.

## Conclusions

In this study, we proposed a solution to the unique challenges encountered by FCS-based methods aimed at probing nuclear mobility without the loss of spatial information. This method is based on slow, continuous line-scanning FCS, that is capable of sampling different nuclear positions while keeping a high temporal resolution. A key aspect of the method is the use of a reference intensity trace to sort segments of the whole FCS measurement into two or more populations corresponding to specific nuclear regions.

We used this method to probe the diffusion of inert GFP in different nuclear domains. As expected, the diffusion was slowed down in the nucleolus relative to the nucleoplasm, presumably due to higher molecular crowding. More interestingly, when studying the GFP mobility in hetero- vs eu-chromatin, we found a slight difference between the diffusion coefficients, which is increased when compared to perinucleolar heterochromatin. A variation of the mobility of GFP was also appreciable when using treatments that alter chromatin structure. These results indicate that, even for a small inert probe like GFP, chromatin and its compaction states can markedly influence diffusion rates, allowing the possibility of using such small inert probes to study chromatin organization and nuclear rheology in living cells. Due to the single-cell sensibility of our method, we were able to highlight intracellular variations of mobility between different chromatin regions, despite a high intercellular variability.

We also showed the applicability of our method in the study of proteins that interacts with chromatin, bringing as an interesting example intranuclear mobility of the Estrogen Receptor. We were able to discriminate between diffusion measured at an engineered prolactin gene array (e.g., transcription locus), where the number of EREs and receptor proteins is higher than in the nucleoplasm. Interestingly, we also retrieved important information about the binding and the residence time of the proteins on the array versus the other binding sites scattered throughout the nucleoplasm.

Finally, we coupled our intensity sorted FCS method with STED microscopy, demonstrating the possibility to probe different nuclear regions at subdiffraction spatial scales. In addition, due to the efficient post-processing tuning of detection volumes, we were able to obtain a spot-variation analysis specific for each compartment in the same measurement.

In summary, we proposed a new, statistically robust method that can be used to perform accurate mobility measurements, at microsecond temporal resolution, in different compartments without the constraint of having the probed regions nearly immobile during FCS measurements. In addition, intensity sorted FCS is suitable to study the diffusion of small inert probes, but also the interaction of proteins with a slowly moving substrate. We believe that this method, especially if coupled with super-resolution, will be useful to study the dynamics of chromatin at the nanoscale, eventually leading to interesting insights into nuclear structure-function relationships.

## Author Contributions

LL, AD and GV designed research. MDB prepared samples and performed experiments. LL wrote software. All authors analyzed data and discussed results. MDB and LL wrote the manuscript with input from AD, GV, DM, MM.

## Acknowledgments

Imaging work was performed at the Nikon Imaging Center @ Istituto Italiano di Tecnologia, generously supported by Nikon. The authors wish to thank Maureen G. Mancini for development of the GFP-ER:PRL-HeLa cell line. LL was supported by Fondazione Cariplo and Associazione Italiana per la Ricerca sul Cancro (AIRC) through Trideo (Transforming Ideas in Oncological Research) Grant ID 17215. DM was supported by Fondazione Cariplo (Ricerca Biomedica condotta da giovani ricercatori, 2014-1157). The authors also wish to thank G. Tortarolo for technical support.

